# Microglia as transcriptional pacemakers of neuronal aging heterogeneity across individuals

**DOI:** 10.64898/2026.04.05.716624

**Authors:** Christine M Lim, Michele Vendruscolo

## Abstract

Neuronal aging pace varies markedly between individuals, but what drives this variation remains unknown. Using cell-type-specific transcriptomic clocks applied to single-nucleus RNA sequencing data from 226 adults (ages 20-90), we quantified neuronal aging residuals as a donor-dominant phenotype. Variance decomposition revealed that microglial transcriptional programs predict inter-individual variation in neuronal aging residuals, a directional asymmetry consistent with a non-cell-autonomous relationship between microglial states and neuronal aging trajectories. This asymmetry is accompanied by an age-dependent shift from homeostatic to inflammatory microglial dominance beginning in midlife, with inflammatory dominance probability rising from 26% at age 35 to 92% by age 65, replicated in an independent cohort. IFNγ signaling emerges as the dominant microglial program associated with accelerated neuronal aging in late adulthood. Candidate regulators of microglial IFNγ activity (*HIF1A, CEBPB*, and *EZH2*) are computationally prioritized as intervention targets warranting functional validation.

## Introduction

Brain aging is a major risk factor for neurodegenerative disease^1-3^, yet individuals of the same chronological age exhibit markedly different rates of molecular and cellular decline^3,4^. Single-cell transcriptomics (sc/snRNA-seq) has enabled the systematic characterization of cell-type-specific transcriptomic changes with age, revealing that aging signatures are largely cell-type-specific and not reducible to a universal program shared across cell types^5,6^. Despite this progress, the factors driving inter-individual variability in neuronal aging pace remain poorly understood^3,4,7^. Extending the logic of epigenetic clocks^8^ to the transcriptome, cell-type-specific transcriptomic aging clocks have emerged as a quantitative framework for measuring this heterogeneity beyond chronological age^4,9^. However, whether the inter-individual variation these clocks reveal is neuron-intrinsic, or shaped by the surrounding cellular environment, remains an open question.

Microglia, the resident immune cells of the brain, are prime candidates for shaping this cellular environment^10,11^. They undergo profound transcriptional state transitions with age, shifting from homeostatic programs toward inflammatory and interferon-responsive states^12,13^. Indeed, this age-associated shift has been reported in aged tissues and neurodegenerative disease contexts^14,15^. Recent spatial transcriptomic atlases have further characterized this relationship, revealing that aging induces spatially dependent cell-state changes^16,17^ and alters microglial proximity to vulnerable neuronal populations^18^. Notably, these spatial shifts are accompanied by pro-aging proximity effects^17,18^. While the non-cell-autonomous effects of microglia on neuronal function are well established in disease^19,20^, it remains unknown whether these microglial state transitions systematically predict inter-individual variation in neuronal aging pace across the adult lifespan^13,16,21,22^.

To investigate whether microglial states predict inter-individual variation in neuronal aging, we integrate cell-type-specific transcriptomic clocks with variance decomposition and in silico perturbation modelling. Aging residuals, which are the deviation of predicted biological age from chronological age^23,24^, provide a donor-level measure of neuronal aging pace. Dominance analysis then quantifies the relative contribution of microglial and neuronal transcriptional programs to these residuals^25^, while perturbation-based control node identification^26-28^ nominates candidate upstream regulators. Together, this framework moves from descriptive association toward directional inference, though it remains subject to the inherent constraints of cross-sectional human data.

We apply this integrated framework to snRNA-seq data from human brains across the adult lifespan (ages 20-90). Cell-type specific transcriptomic aging clocks were constructed and specific microglial programs capable of predicting deviations in neuronal aging were identified. Our analysis reveals a directional asymmetry, as microglial states predict neuronal aging pace. We demonstrate that this interaction is accompanied by an age-dependent shift from homeostatic to inflammatory microglial dominance beginning midlife. IFNγ signaling emerges as the primary microglial axis associated with neuronal aging in late adulthood, with upstream transcription factor control nodes nominated for future functional validation. Together, this study proposes a non-cell-autonomous framework to map the regulatory control nodes that govern inter-individual variation in aging rates, providing a data-driven foundation for precision geroscience.

## Results

### Excitatory neurons encode a reproducible transcriptomic aging signal captured by cell-type-specific clocks

To characterize cell-type-specific aging dynamics in the human brain, we constructed transcriptomic clocks using donor-level pseudobulk snRNA-seq profiles across the adult lifespan (ages 20-90; **Fig. 1A**). We trained elastic net regression models for eight major brain cell types **(Fig. 1B)**, among which the excitatory neuronal model achieved strong predictive performance (MAE = 6.0 years, Pearson r = 0.90, 5-fold cross-validation). This performance was replicated in an independent external validation dataset **(Fig. 1C)**, indicating that excitatory neuron transcriptomes encode a robust and conserved molecular aging signal.

**Figure 1.**
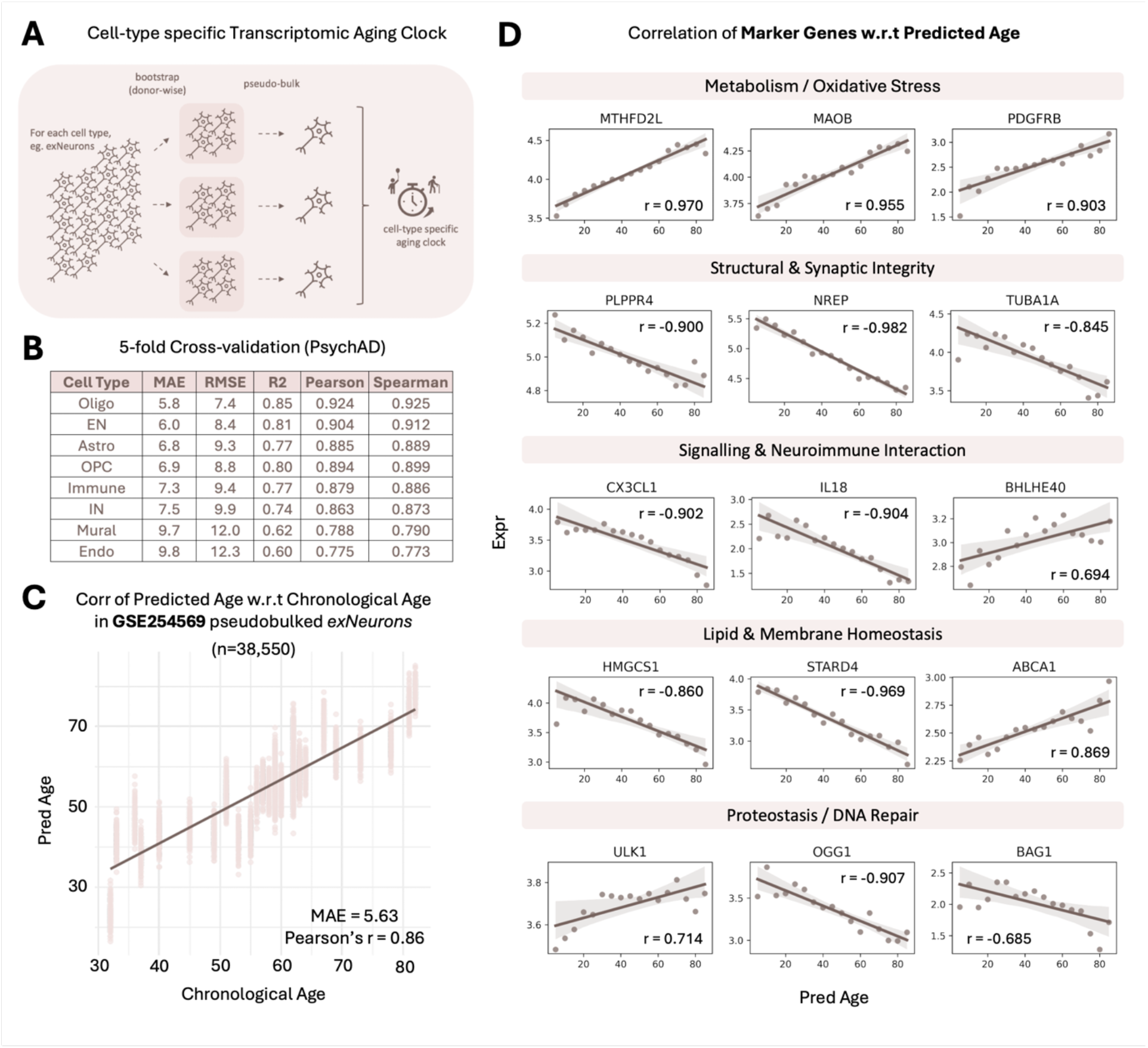
Construction and validation of cell-type-specific transcriptomic aging clocks identify a robust aging program in excitatory neurons. **(A)** Schematic of cell-type-specific donor-wise pseudobulking procedure. For each donor, cells of the target cell type were repeatedly sampled with replacement across 150 bootstrap iterations and averaged to generate donor-level pseudobulk expression profiles. **(B)** Cross-validation performance (5-fold) of cell-type-specific transcriptomic aging clocks. **(C)** Predicted biological age versus chronological age for excitatory neurons in GSE254569 (external dataset). Each point represents a single donor-level pseudobulk profile averaged across bootstrap replicates. The solid line shows the best-fit linear regression. **(D)** Mean expression of 15 monotonically aging marker genes across predicted age bins. Genes are organized into five functional modules: metabolism/oxidative stress (*MTHFD2L, MAOB*, and *PDGFRB*), structural and synaptic integrity (*PLPPR4, NREP*, and *TUBA1A*), neuroimmune signaling (*CX3CL1, IL18*, and *BHLHE40*), lipid and membrane homeostasis (*HMGCS1, STARD4*, and *ABCA1*), and protein quality control/DNA repair (*ULK1, ATG5, BAG1*). Each point represents mean expression within a 5-year predicted-age bin. Solid lines show linear best-fit trends with 95% confidence intervals. Spearman correlation coefficients (r) are reported per gene.

To ensure the biological validity of this signal, we observed 15 aging marker genes that organized into five functional modules: metabolism/oxidative stress, structural and synaptic integrity, neuroimmune signaling, lipid and membrane homeostasis, and protein quality control/DNA repair **(Fig. 1D)**. We found that the aging markers exhibited monotonic associations with predicted age (|r| > 0.6, FDR < 0.05). To confirm that these markers represent a non-random biological signature, we compared their mean absolute correlations against two null distributions: one generated from 1,000 bootstrapped gene sets (n=15) and another generated from shuffled age labels. The selected markers significantly outperformed both null models **(Fig. S1)**, confirming that the clock captures a coordinated biological aging program.

### Neuronal aging pace is a donor-specific phenotype

Having established a robust neuronal aging clock, we next investigated the inter-donor heterogeneity in biological aging pace. While predicted neuronal age correlated with chronological age, we observed substantial variability within age bins, revealing that individuals of similar chronological age can exhibit markedly different molecular aging profiles **(Fig. 2A)**. We hence quantified this divergence using age-adjusted residuals, i.e. the deviation of each donor’s predicted neuronal age from their chronological age **(Fig. 2B)**.

**Figure 2.**
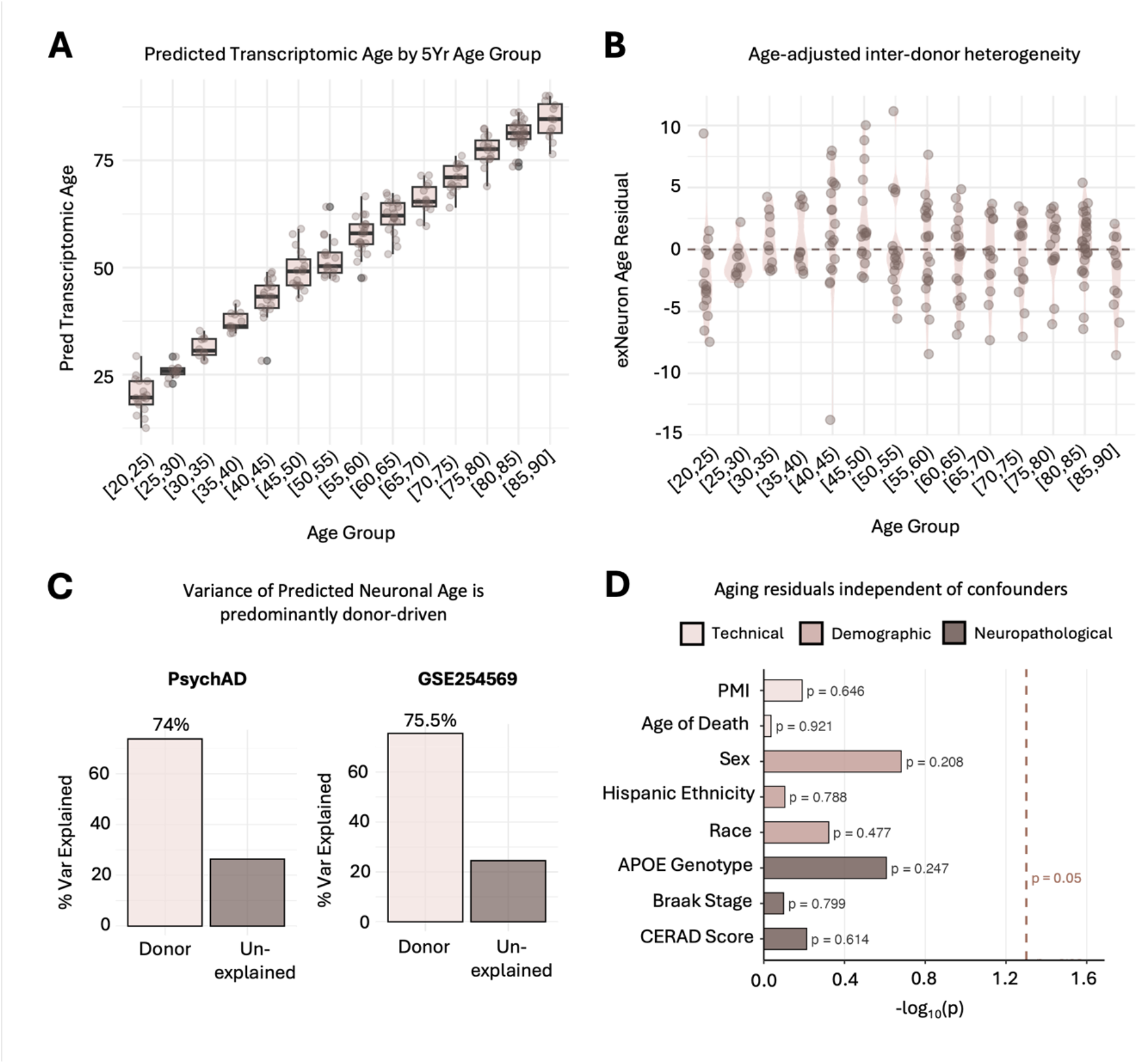
Neuronal aging residuals are a robust donor-specific phenotype independent of technical and neuropathological confounders. **(A)** Predicted neuronal age by chronological age bin. Significant inter-donor variability within 5-year bins reveals that molecular aging pace diverges meaningfully between age-matched individuals. **(B)** Age-adjusted neuronal aging residuals. Violin plots show the deviation from chronological age (dashed line = 0) across adulthood. **(C)** Variance architecture of transcriptomic aging. Linear mixed-effects modeling shows donor identity explains the vast majority of variance in predicted age (74% in the PsychAD dataset and 75.5% in GSE254569), confirming that aging pace is a robust donor-specific feature. **(D)** Confounder analysis. Associations between neuronal aging residuals and eight technical, demographic, and neuropathological variables. All associations are non-significant (p > 0.05 for all variables; dashed line indicates significance threshold). Full distributions for each confounder are reported in **Figure S2**.

Variance decomposition revealed that donor identity accounts for 74% of the variance in predicted neuronal age **(Fig. 2C)**. This dominant contribution of donor identity to the total variance was replicated in an independent cohort (GSE254569^29^; 75.5%; **Fig. 2C**), confirming that inter-individual differences in neuronal aging pace are a robust biological phenomenon rather than a dataset-specific artifact.

To assess whether the age-adjusted residuals reflect genuine biological variation rather than technical or demographic confounds, we tested for associations with post-mortem interval, sex, ethnicity, race, and APOE genotype, finding no significant correlations **(Figs. 2D and S2)**. Notably, among donors with available neuropathological data (n = 139), aging residuals showed no association with Braak stage^30^ or CERAD score^31^ **(Figs. 2D and S2)**. This suggests that the excitatory neuron clock captures molecular aging variation that is statistically dissociable from classical neuropathological staging in this cross-sectional cohort, though whether this signal reflects processes that are biologically independent of, or temporally precede, aggregate accumulation cannot be resolved without longitudinal data.

### An age-dependent shift in microglial program dominance is associated with asymmetric prediction of neuronal aging residuals

To identify the transcriptional drivers of the neuronal aging offset, we evaluated the relative contribution of microglial and neuronal hallmark programs **(Methods)**. Collectively, we find that microglial and neuronal hallmark programs explain ∼22% of neuronal age residual variance **(Fig. 3A)**. Glycolysis, DNA repair, and the P53 pathway were neuronal transcriptomic programs that best explained neuronal aging residual variance, while IFNγ response, TNFα signalling, and oxidative phosphorylation were microglial programs that best explained neuronal residual variance.

**Figure 3.**
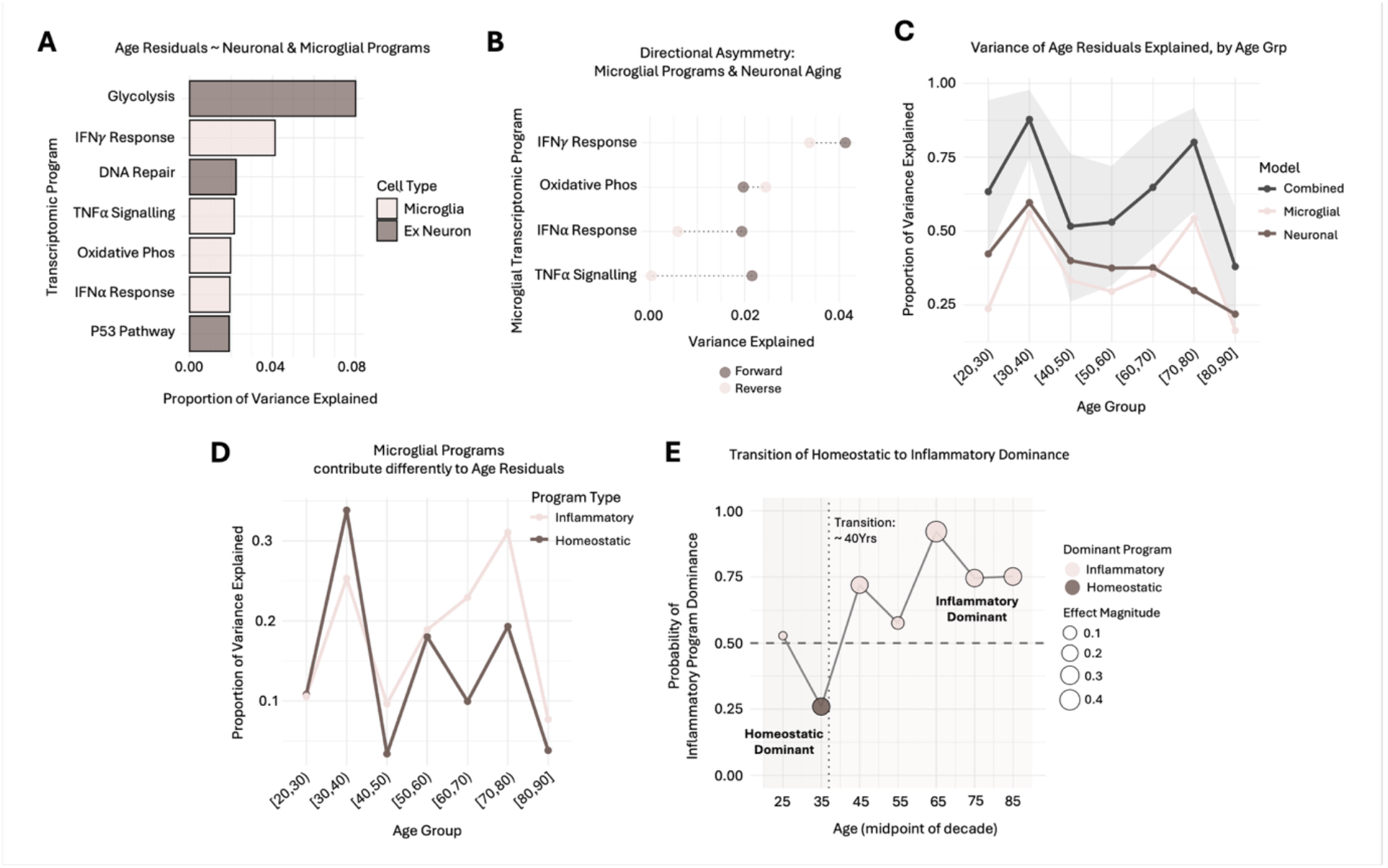
Microglial programs asymmetrically predict neuronal aging residuals and undergo an age-dependent homeostatic-to-inflammatory transition. **(A)** Transcriptomic program variance decomposition. Relative importance of microglial and neuronal hallmark programs in predicting neuronal aging residuals. Specific immune and homeostatic programs emerge as top predictors, collectively explaining ∼22% of residual variance. **(B)** Comparison of forward versus reverse variance explanation reveals that microglial states are associated with neuronal aging pace. **(C)** Combined microglial-neuronal models consistently outperform cell-type-specific models across adulthood. Notably, a dual peak in microglial programs influence on neuronal aging pace is observed in early and late adulthood. Shaded ribbon indicates 95% bootstrap CI. **(D)** Contribution of microglial functional axes across adulthood. Homeostatic programs are the primary drivers of neuronal residuals in early adulthood, while inflammatory programs emerge as the dominant predictive axis from midlife onward. **(E)** Quantification of the homeostatic-to-inflammatory shift. The probability of inflammatory dominance was quantified and clearly reflects the age range at which inflammatory programs outweigh homeostatic programs. Dominance rises sharply after age 40, peaking at 92% in late adulthood.

To test the directionality of this relationship, we compared the variance explained by microglial programs on neuronal residuals (Forward) against the variance explained by neuronal residuals on those same microglial programs (Reverse). We observed a directional asymmetry: microglial programs explained substantially more variance in neuronal aging residuals than the reverse **(Fig. 3B)**. This statistical lead indicates that microglial state is predictive of neuronal aging pace, consistent with a model in which microglial states contribute to neuronal aging pace non-cell-autonomously, although this asymmetry could also arise from microglia being more transcriptionally variable across donors or from confounding by shared upstream factors such as systemic inflammation or vascular health^32^.

We next examined how these contributions evolve across the adult lifespan. Across all age bins, a combined model of microglial and neuronal programs consistently explained more than 25% of the variance in neuronal residuals **(Fig. 3C)**. Interestingly, we observed two peaks in the contribution of microglial programs toward explaining neuronal aging residual variances in early adulthood vs late adulthood. Decomposing these microglial contributions by functional types revealed a systematic, age-dependent shift in the drivers of this asymmetry **(Fig. 3D)**. While homeostatic microglial programs were the primary contributors in early adulthood, inflammatory programs progressively emerged as the dominant axis from midlife onward. To quantify this transition, we computed the probability of inflammatory dominance across 500 bootstrap replicates per age bin **(Fig. 3E)**. Inflammatory dominance probability was low in early adulthood (26% at ages 30-40) but rose sharply from midlife.

### Microglial IFNγ response is the dominant predictor of neuronal aging residuals, with candidate upstream regulators identified by network modelling

To identify which microglial processes exert the greatest influence on the neuronal aging offset, we performed perturbation simulations within a joint linear model of neuronal and microglial programs (Methods). By conducting scenario modelling within the joint linear model, varying each program’s activity score independently, we could estimate the predicted contribution of each microglial program to donor neuronal aging residuals within the joint linear model. Among all tested programs, microglial IFNγ response was associated with the largest positive shift in predicted neuronal aging residuals across the donor cohort **(Fig. 4A)**. Bootstrapped confidence intervals confirmed the statistical robustness of this estimate across resampled donor subsets, ranking microglial IFNγ response as the largest positive predictor of neuronal aging residuals within the tested set of HALLMARK microglial program^27^, although this ranking is necessarily constrained by the programs selected for analysis.

**Figure 4.**
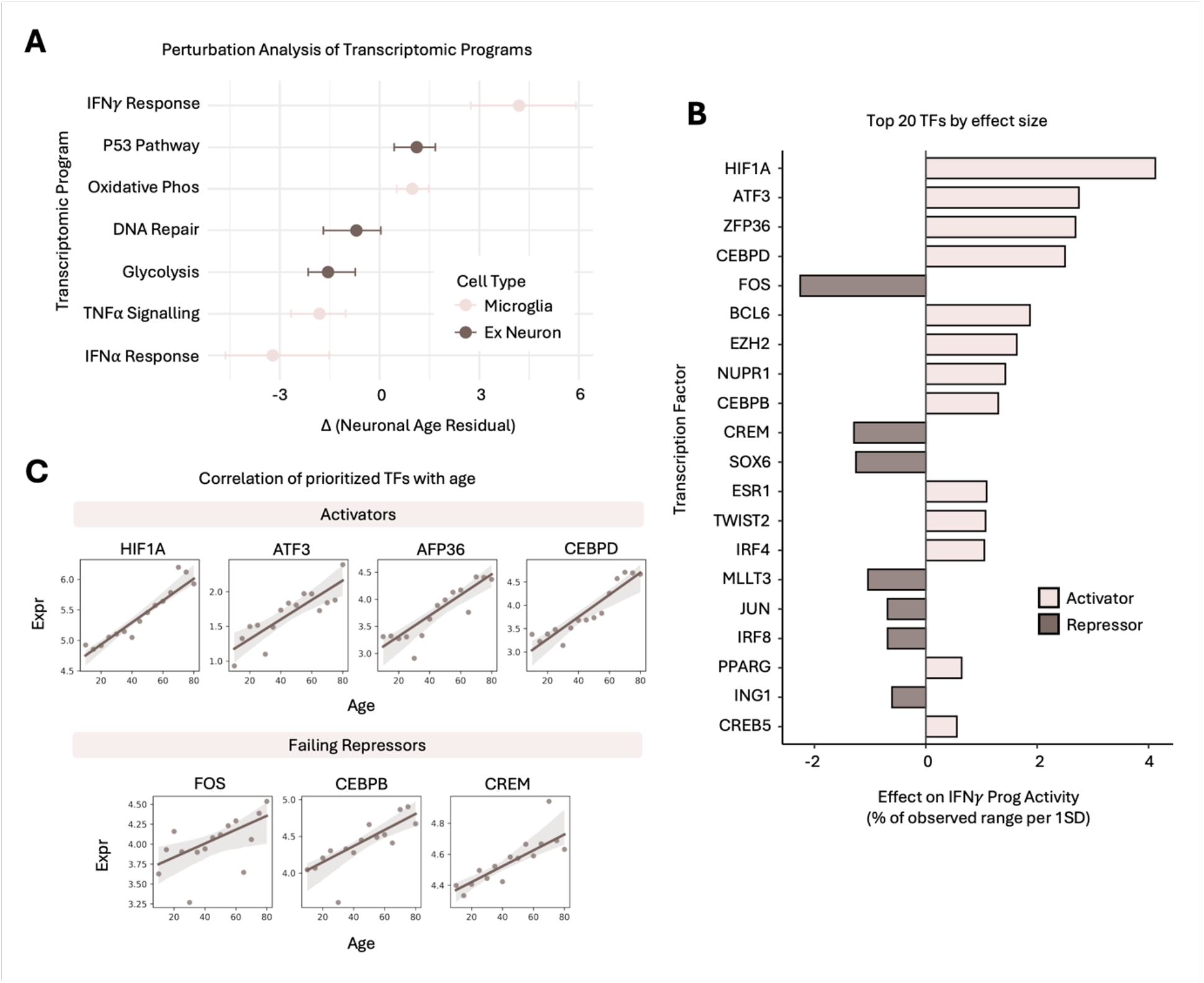
IFNγ response is the dominant microglial program predicting neuronal aging residuals and is associated with a prioritized transcription factor network. **(A)** In silico perturbations of aging programs and their predicted effects on neuronal aging residuals. Microglial IFNγ response produces the largest positive shift in neuronal age residuals among all tested programs. Error bars represent 95% bootstrap CIs. **(B)** Transcriptional control nodes of microglial IFNγ activity: the top 20 transcription factors by eiect size are presented. Coeiicients represent the percentage of the observed IFNγ range shifted per 1 SD change in TF expression. Top putative activators include *HIF1A, ATF3*, and *ZFP36;* while *FOS* and *CREM* are putative repressors. **(C)** Repressor paradox of microglial aging: while activators show expected age-associated increases, identified putative repressors also exhibit positive correlations with age. This concurrent upregulation of putative repressors alongside escalating IFNγ response activity suggests a state of high-regulatory tension and the impairment of endogenous homeostatic buffering in aging microglia.

To identify the transcriptional control nodes of IFNγ response, we prioritized transcription factors that best predict IFNγ program activity scores (Methods). The top activators by effect size included *HIF1A, ATF3*, and *ZFP36* **(Fig. 4B)**. Among the prioritized regulators, HIF1A is notable because it lies within a pharmacologically tractable hypoxia-inducible factor (HIF) signaling pathway^33,34^. However, the convergence of a computationally nominated regulator with an existing drug target, while hypothesis-generating, does not constitute independent validation of either the regulatory relationship or the relevance of metformin to the microglial IFNγ axis described here^35^.

We quantified the correlation of the top TFs with donor age (**Fig. 4C**), confirming that activator expression increases with age as expected. Intriguingly, the putative repressors *FOS, CEBPB*, and *CREM*, identified as negative regulators of IFNγ response activity, also showed positive correlations with age, despite their inhibitory classification. This paradoxical co-occurrence is consistent with escalating regulatory tension, though it could also reflect co-expression across transcriptionally distinct microglial subpopulations that is obscured by pseudobulk averaging.

We interpret this pattern as consistent with insufficient endogenous buffering. The age-associated upregulation of FOS likely reflects a compensatory attempt by the AP-1 resolution pathway to restrain escalating neuroinflammation. This interpretation is supported by literature on AP-1 as a context-dependent repressor of interferon signalling^36,37^. Similarly, the parallel upregulation of *CEBPB* and *CREM* may represent a coordinated but ultimately insufficient homeostatic response. Critically, the co-expression of activators and repressors does not appear to produce equilibrium: IFNγ program activity continues to rise with age (**Fig. 4C**), suggesting that repressor upregulation is quantitatively outpaced by the dominant inflammatory drive.

This pattern closely mirrors regulatory dynamics observed in microglial senescence, wherein pro-inflammatory programs and their nominal inhibitors are simultaneously elevated in a state that resists resolution^38,39^. Importantly, if endogenous repressors are already engaged but insufficient, therapeutic strategies aimed solely at boosting inhibitory signalling may be limited in efficacy. Distinguishing whether repressor insufficiency reflects quantitative outcompetition, epigenetic decoupling from target loci, or expression in transcriptionally distinct microglial subpopulations remains an important question for future work, requiring chromatin accessibility or single-cell TF binding data beyond the scope of this study.

## Discussion

In this study, we have shown that the neuronal aging pace, quantified as the residual deviation between transcriptomic-predicted and chronological ages, is associated with microglial transcriptional states in a directionally asymmetric manner. By integrating cell-type-specific clocks, variance decomposition, and perturbation simulations across the adult human lifespan, we have reported three main findings: First, microglial programs predict neuronal aging residuals, supporting a model of non-cell-autonomous regulation. Second, the microglial programs underlying this asymmetry undergo a systematic age-dependent shift from homeostatic dominance in early adulthood to inflammatory dominance starting in midlife. Third, IFNγ signaling emerges as a dominant microglial axis associated with accelerated neuronal aging in late adulthood, regulated by a prioritized set of transcription factors.

The directional asymmetry that we observed builds upon existing literature reporting non-cell-autonomous effects of microglia on neuronal integrity in disease, extending the principle to the normal aging trajectory^22,40,41^. The homeostatic-to-inflammatory shift in microglial dominance, beginning in midlife and consolidating in late adulthood, is consistent with emerging evidence of microglial homeostatic reprogramming^13,21,42^. By quantifying the probability of this transition, we have identified a midlife period during which inflammatory dominance is consolidating but not yet saturated. Whether this period represents a window in which microglial trajectories are more amenable to redirection will depend on establishing causality through longitudinal and interventional studies. More broadly, the age-associated co-upregulation of putative IFNγ activators and repressors such as FOS suggests that endogenous buffering mechanisms may be engaged but insufficient to counterbalance the dominant pro-inflammatory drive^12,36^. Defining the conditions under which such regulatory restraints fail may help clarify how microglial homeostatic control is progressively lost during aging^11,13^.

Several limitations should be considered when interpreting these findings. First, this study is based on cross-sectional post-mortem data, which constrains our ability to establish temporal ordering or causal directionality. The observed directional asymmetry is consistent with a non-cell-autonomous model but cannot exclude alternative explanations: microglia may exhibit greater inter-individual transcriptional variability than neurons, making them statistically stronger predictors of any donor-level phenotype, or both cell types may respond to shared upstream factors such as systemic inflammation, vascular dysfunction, or peripheral immune signalling that are not captured in this dataset^43^. Resolving whether microglial states causally influence neuronal aging pace, or merely co-vary with it, will require longitudinal sampling and experimental perturbation in model systems. Second, all analyses were performed on pseudobulk profiles, which average across transcriptionally distinct microglial subpopulations within each donor. The homeostatic-to-inflammatory transition that we described could reflect genuine within-cell reprogramming, but it could equally arise from shifting proportions of pre-existing microglial subpopulations^44^. This is a distinction that could be resolved by single-cell-resolution analyses or spatial deconvolution approaches. The repressor paradox, in which putative inhibitors of IFNγ activity are co-upregulated with activators, is particularly susceptible to this confound, as co-expression at the pseudobulk level may reflect distinct subpopulations expressing activators and repressors separately. Third, the discovery cohort is derived from prefrontal cortex, and microglial transcriptional states are known to vary substantially across brain regions^45^. Whether the microglial programs identified here, and the midlife transition in their dominance, generalise to hippocampus, white matter, or subcortical regions remains to be determined. Fourth, the transcription factor control nodes (*HIF1A, EZH2*, and others) were nominated computationally through regularised regression on curated TF-target databases, without experimental validation by chromatin accessibility profiling, perturbation experiments, or independent cohort replication of the regulatory relationships. Although HIF1A lies within a pharmacologically tractable signalling pathway, this convergence does not constitute independent validation of the inferred regulatory relationship or the relevance of any specific agent to the microglial IFNγ axis identified here^33^. These nominations should therefore be regarded as hypothesis-generating rather than as validated regulatory mechanisms. Finally, while we tested for associations between neuronal aging residuals and available demographic, technical, and neuropathological variables, unmeasured confounders, including medication history, comorbidities, lifetime inflammatory burden, and cause of death, could contribute to the inter-individual variation attributed here to microglial transcriptional states.

Overall, the computational framework developed here integrates transcriptomic clocks, variance decomposition, and perturbation modelling to generate a prioritised set of candidate targets and a provisional timeline for the microglial homeostatic-to-inflammatory transition. Whether this framework can inform geroprotective strategy will depend on longitudinal replication of the transition dynamics, experimental validation of the nominated regulatory nodes, and demonstration that modulating these nodes alters neuronal aging outcomes. The most immediate next steps include longitudinal sampling to confirm the midlife transition within individuals rather than across cohorts, experimental perturbation of the prioritised transcription factors in microglial culture or mouse models, and extension of the variance decomposition to additional brain regions and cell types to assess the generality of the microglia-excitatory neuron asymmetry. In conclusion, this work illustrates how integrating transcriptomic clocks with variance decomposition and perturbation modelling can nominate candidate cellular programs and regulatory nodes for further investigation, a necessary step toward determining whether such programs are experimentally modifiable and whether their modulation would alter aging trajectories in vivo.

## Methods

### Data acquisition and cohort integration

The discovery cohort was constructed by integrating snRNA-seq data from the PsychAD consortium (aging and RADC cohorts)^46,47^. To characterize non-pathological aging, the unified dataset was restricted to cognitively healthy donors aged ≥ 20 years. External validation was performed using the GSE254569^29^ dataset, an independent cohort of non-overlapping donors.

### Quality control and cell-type stratification

Count matrices were filtered to retain protein-coding genes only. Cell-level quality control excluded nuclei expressing fewer than 200 or more than 6,000 genes; genes detected in fewer than 10 cells were also removed^48^. Analyses were stratified by cell type, with excitatory neurons serving as the primary focus for transcriptomic clock construction and microglia for functional program scoring and regulatory node inference.

### Cell-type-specific pseudobulking

To robustly capture donor-level signal while accounting for single-cell sparsity, we employed a bootstrap pseudobulking procedure^9^. For each donor, cells of the target type were normalized to counts per million (CPM). We then generated 150 bootstrap iterations per donor by randomly sampling 50 cells (microglia, astrocytes, OPCs, immune cells, mural cells, and endothelial cells); 100 cells (excitatory and inhibitory neurons); and 200 cells (oligodendrocytes) with replacement^9^ and computing the mean normalized expression vector. Pseudobulk profiles were generated independently for each cell type.

### Transcriptomic clock construction and validation

Cell-type specific aging clocks were trained using Elastic Net regression (ElasticNetCV; Python) on pseudobulk cells^49^. The top 3000 HVGs were z-score standardized and used for training the aging clock. To mitigate age-group imbalance, we applied inverse-frequency sample weights based on 10-year bins during training. All pseudobulk cells from a single donor were assigned exclusively to either training or held-out sets. To correct for systematic age-dependent prediction drift, we applied linear proportional-fold correction (L-PFC)^4,9^. Model performance was quantified using MAE, R^2^, Pearson, and Spearman correlations on held-out folds. The final model for downstream inference was trained on the complete discovery cohort using this optimized pipeline.

### Neuronal aging residual derivation and variance decomposition

To quantify inter-individual biological variation of neuronal aging, we derived neuronal aging residuals. Predicted ages from bootstrap replicates were first averaged to produce a single donor-level estimate. The donor-level predicted ages were then regressed against chronological age. The resulting residuals were mean-centered and utilized for all downstream regulatory analyses. To quantify the stability of this signal, we performed variance decomposition using a linear mixed-effects model (lme4; R)^50^: predicted neuronal age was modeled with chronological age as a fixed effect and donor identity as a random intercept. Variance components were extracted via restricted maximum likelihood (REML) and expressed as a percentage of total variance to determine the proportion of aging pace attributable to stable donor-specific factors versus stochastic or technical noise.

### Confounder analysis

To assess the association between neuronal aging residuals and potential confounders: Spearman rank correlation was used for continuous variables (post-mortem interval, age of death). Wilcoxon rank-sum tests were used for binary categorical variables (sex, Hispanic ethnicity, CERAD score grouped into ‘none’ vs. ‘any amyloid’). Kruskal-Wallis tests were used for variables with three or more groups (race, APOE genotype, Braak stage). All available data were used for these analyses.

### Transcriptional program scoring

Cell-type-specific program activities were quantified using AUCell^51^ applied to pseudobulked expression profiles. Neuronal programs were defined using curated HALLMARK gene sets from MSigDB^27^: GLYCOLYSIS, P53_PATHWAY, DNA_REPAIR, MTORC1_SIGNALING, NOTCH_SIGNALING, OXIDATIVE_PHOSPHORYLATION, PEROXISOME, REACTIVE_OXYGEN_SPECIES_PATHWAY, UNFOLDED_PROTEIN_RESPONSE, and WNT_BETA_CATENIN_SIGNALING. Microglial programs were also defined using relevant HALLMARK gene sets^27^: APOPTOSIS, COMPLEMENT, GLYCOLYSIS, IL6_JAK_STAT3_SIGNALING, INFLAMMATORY_RESPONSE, INTERFERON_ALPHA_RESPONSE, INTERFERON_GAMMA_RESPONSE, OXIDATIVE_PHOSPHORYLATION, PROTEIN_SECRETION, and TNFA_SIGNALING_VIA_NFKB.

For downstream analyses, influential microglial programs were classified into two functional categories: homeostatic programs (INTERFERON_ALPHA_RESPONSE and OXIDATIVE_PHOSPHORYLATION) and inflammatory programs (INTERFERON_GAMMA_RESPONSE and TNFA_SIGNALING_VIA_NFKB).

### Relative importance and directional asymmetry analysis

To quantify the contribution of each curated transcriptomic program to the neuronal aging pace, we fit a joint linear regression model using standardized AUCell scores as predictors of neuronal aging residuals. The relative importance of each program was estimated using LMG dominance analysis (relaimpo; R)^52^. Directional asymmetry was assessed by comparing forward variance explained (microglial program > neuronal aging residual) against the reverse. To examine temporal shifts in these relationships, donors were stratified into 10-year age bins. Within each bin, bootstrapping was used (100 replicates, sampling donors with replacement) to compute the mean R^2^ and 95% confidence intervals for neuronal-only, microglial-only, and combined predictor sets.

### Bootstrap dominance probability analysis

To quantify the transition in microglial program influence, we computed the probability of inflammatory dominance across 500 bootstrap replicates per 10-year age bin. For each replicate, dominance was defined as the inflammatory program set explaining relatively more variance (R^2^) in neuronal residuals than the homeostatic set.

### In silico perturbation and regulatory network modelling

We carried out in silico perturbation simulations to estimate the model-predicted directional effect of transcriptional programs on neuronal residuals. Within the joint linear model framework, each program was independently shifted by +2 standard deviations while holding other predictors at observed values. The resulting mean shift in predicted neuronal aging residual (Delta Y) was averaged across all donors and bootstrapped over 500 iterations to generate 95% confidence intervals.

To identify upstream regulators of the dominant IFNγ response program, we curated a transcription factor (TF)-target network from TRRUST^53^, DoRothEA^54^, and ChEA^54,55^. We filtered for TFs with documented regulatory links to the INTERFERON_GAMMA_RESPONSE hallmark set and extracted their standardized expression from microglial pseudobulk profiles. An Elastic Net regression model (cv.glmnet; R)^56^ was utilized to predict donor-level IFNγ response program activity scores from TF expression. TFs with non-zero coefficients were classified as activators or repressors, and their effect sizes were normalized to the percentage of the observed IFNγ activity score range.

## Data Availability

The single-nucleus RNA sequencing data used in this study are publicly available. The PsychAD consortium data (aging and RADC cohorts) are available from https://psych-ad.org/. The external validation dataset is available from the Gene Expression Omnibus under accession GSE254569.

## Code Availability

All code for transcriptomic clock construction, variance decomposition, dominance analysis, in silico perturbation simulations, and transcription factor network modelling is available at [the GitHub URL will be made available upon publication].

## Author Contributions

C.M.L. and M.V. conceived and designed the study. C.M.L. performed all computational analyses, including clock construction, variance decomposition, perturbation modelling, and regulatory network inference. C.M.L. and M.V. interpreted the results and wrote the manuscript. Both authors reviewed and approved the final version.

## Competing Interests

The authors declare no competing interests.

## Acknowledgements

Data were obtained from the PsychAD consortium.

## Funding

This work was supported by UKRI (10059436, 10061100 and 10138075).

## Ethics Statement

This study used de-identified post-mortem human brain tissue samples obtained from publicly available datasets. No additional ethical approval was required for the secondary computational analyses performed in this study.

## SUPPLEMENTARY INFORMATION

**Figure S1.**
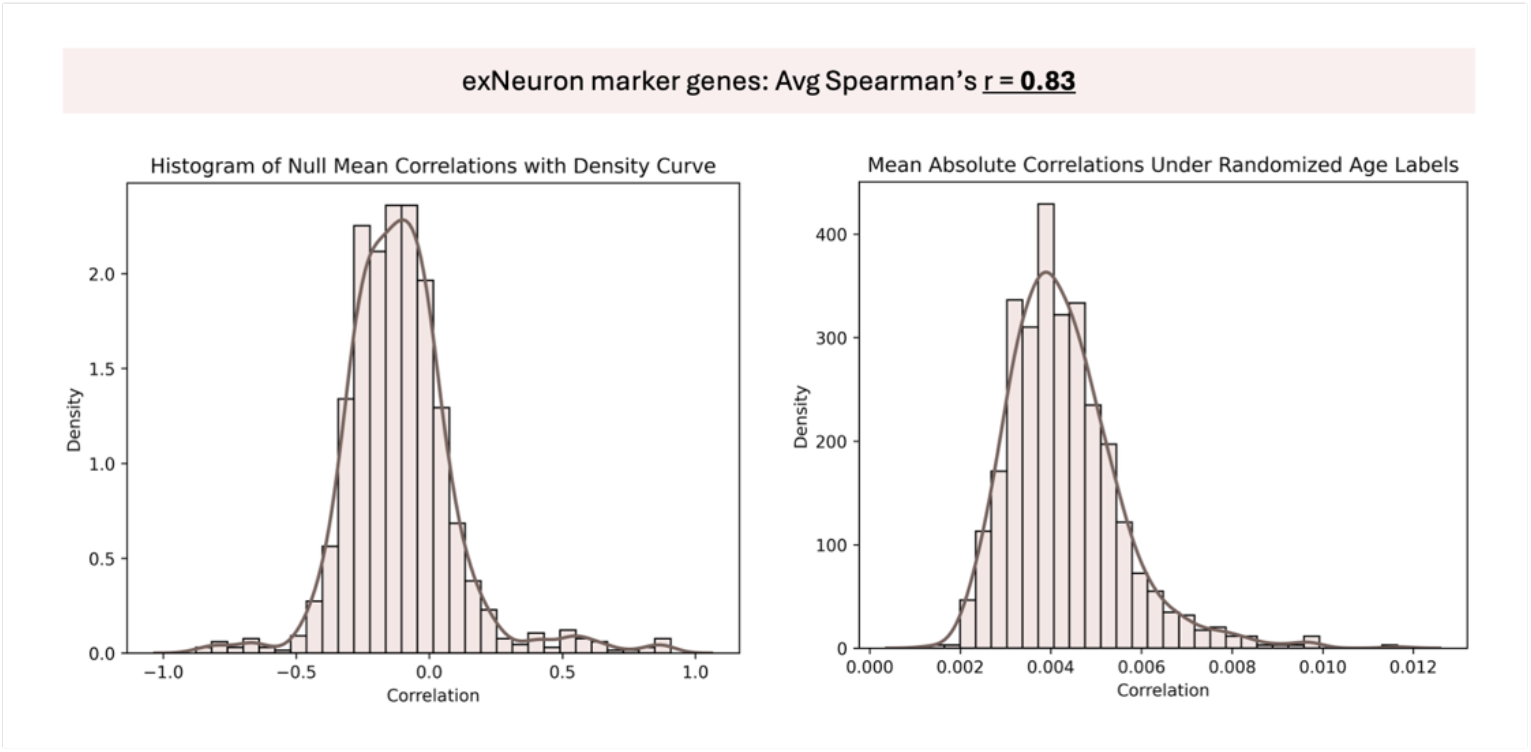
Validation of the neuronal aging marker genes against null models. Performance of aging marker genes against bootstrapped null distributions. The mean absolute correlation of the 15 selected aging markers with predicted age (Spearman r = 0.83) is significantly higher than that of 1,000 randomly sampled gene sets of the same size. Permutation testing of age associations was conducted by comparing the neuronal aging marker-age correlations against a null distribution generated by 1,000 iterations of shuffled chronological age labels. The selected markers consistently outperform the shuffled null models, demonstrating that the clock-age association is driven by genuine age-related transcriptomic variance.

**Figure S2.**
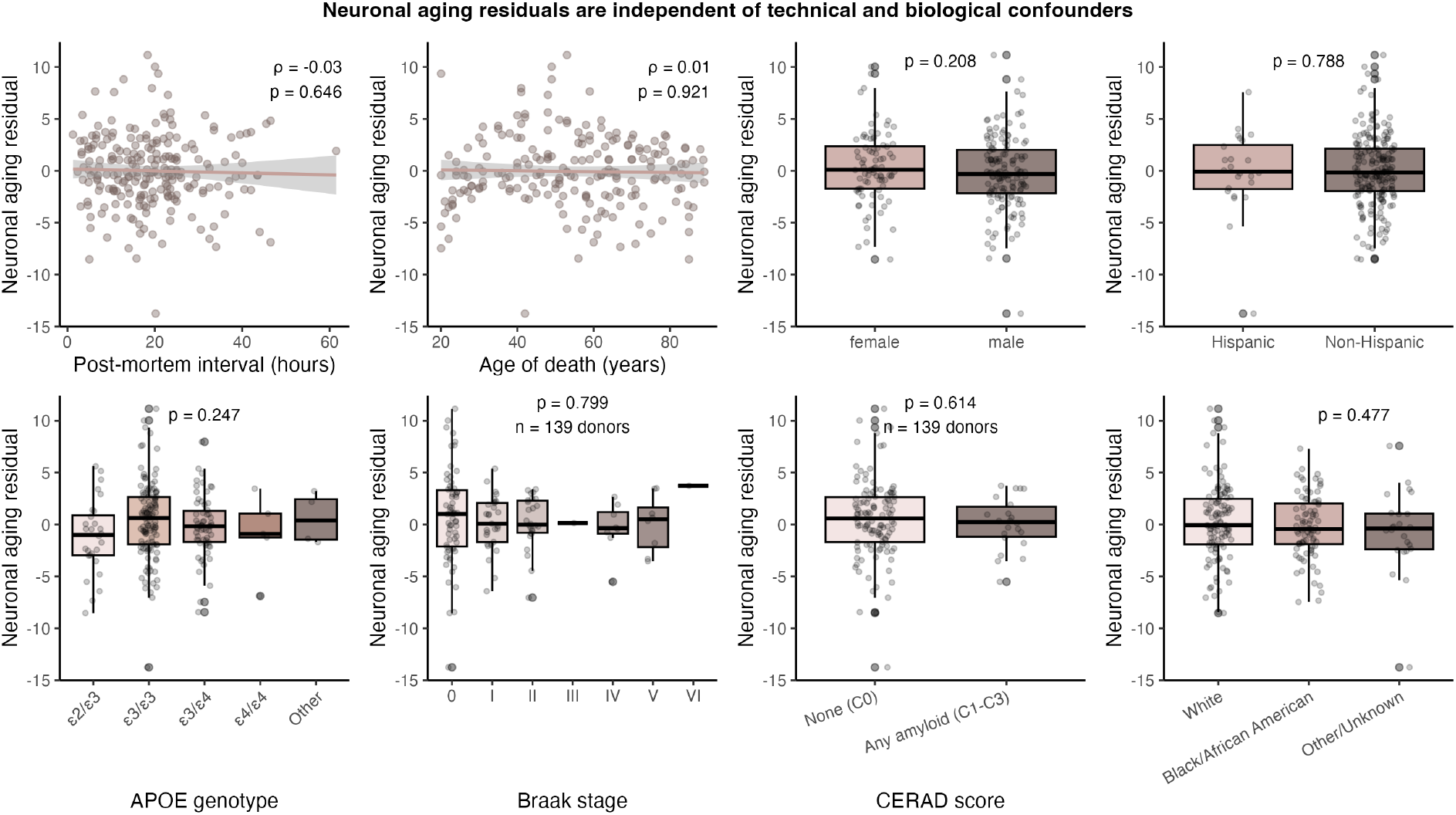
Neuronal aging residuals are independent of technical, demographic, and neuropathological confounders. Residuals are independent of the following: (1) Technical and demographic confounders - post-mortem interval (PMI) or age of death. Scatter plots include linear regression fits with 95% CIs; (2) biological and demographic strata - sex, Hispanic ethnicity, race, and APOE genotype. Boxplots represent the median and IQR, with individual donor points jittered; (3) neuropathological annotations - Braak stage or CERAD score. All associations are non-significant (p > 0.05). Statistical tests used include Spearman rank correlation (technical and demographic confounders), Wilcoxon rank-sum test (biological and demographic strata), and Kruskal-Wallis test (neuropathological annotations).

## Notes

### Competing Interest Statement

The authors have declared no competing interest.

